# Genetic and environmental circadian disruption induce metabolic impairment through changes in the gut microbiome

**DOI:** 10.1101/2022.07.27.501612

**Authors:** Baraa Altaha, Marjolein Heddes, Violetta Pilorz, Yunhui Niu, Elizaveta Gorbunova, Michael Gigl, Karin Kleigrewe, Henrik Oster, Dirk Haller, Silke Kiessling

## Abstract

**Objective:** Internal clocks time behavior and physiology, including the gut microbiome in a circadian (∼24 h) manner. Mismatch between internal and external time, e.g. during shift work, disrupts circadian system coordination promoting the development of obesity and type 2 diabetes (T2D). Conversely, body weight changes induce microbiota dysbiosis. The relationship between circadian disruption and microbiota dysbiosis in metabolic diseases, however, remains largely unknown.

**Methods:** Core and accessory clock gene expression in different gastrointestinal (GI) tissues were determined by qPCR in two different models of circadian disruption - mice with Bmal1 deficiency in the circadian pacemaker, the suprachiasmatic nucleus (*Bmal1^SCNfl/-^*), and wild-type mice exposed to simulated shift work (SSW). Body composition and energy balance were evaluated by nuclear magnetic resonance (NMR), bomb calorimetry, food intake and running-wheel activity. Intestinal permeability was measured in an Ussing chamber. Microbiota composition and functionality were evaluated by 16S rRNA gene amplicon sequencing, PICRUST2.0 analysis and targeted metabolomics. Finally, microbiota transfer was conducted to evaluate the functional impact of SSW-associated microbiota on the host’s physiology.

**Results:** Both chronodisruption models show desynchronization within and between peripheral clocks in GI tissues and reduced microbial rhythmicity, in particular in taxa involved in short-chain fatty acid (SCFA) fermentation and lipid metabolism. In Bmal1SCNfl/- mice, loss of rhythmicity in microbial functioning associates with previously shown increased body weight, dysfunctional glucose homeostasis and adiposity. Similarly, we observe an increase in body weight in SSW mice. Germ-free colonization experiments with SSW- associated microbiota mechanistically link body weight gain to microbial changes. Moreover, alterations in expression of peripheral clock genes as well as clock-controlled genes (CCGs) relevant for metabolic functioning of the host were observed in recipients, indicating a bidirectional relationship between microbiota rhythmicity and peripheral clock regulation.

**Conclusions:** Collectively, our data suggest that loss of rhythmicity in bacteria taxa and their products, which likely originates in desynchronization of intestinal clocks, promotes metabolic abnormalities during shift work.

## 1 Introduction

Most species have evolved endogenous circadian clocks to facilitate adaption to daily recurring changes. A complex hierarchical circadian system consists of a central clock in the suprachiasmatic nuclei (SCN) of the hypothalamus which regulates rhythmic behavior, such as rest-activity, and synchronizes peripheral clocks via neuronal and humoral signals to adapt to environmental changes [1]. Peripheral circadian clocks have been identified in various organs, including the gastrointestinal (GI) tract, and regulate tissue-specific functions, such as glucocorticoid synthesis and glucose metabolism [2; 3]. On the molecular level, the circadian clock consists of a subset of interconnected clock genes which regulate circadian rhythms of tissue-specific clock-controlled genes (CCGs) and thereby control various aspects of physiology [4].

Mismatch between the internal clock and the environmental time, observed in shift workers, induces circadian desynchronization among peripheral clocks [5]. Genetically and environmentally induced circadian disruption has been associated with various metabolic and GI diseases including obesity and diabetes [6–8]. Similarly, lack of the coordinative input from the central clock results in desynchronization between peripheral clocks and causes an increase in body weight and impaired glucose tolerance [6; 9]. These results suggest that peripheral circadian desynchronization might be causal for metabolic alterations.

In the context of metabolic disease, human cohort studies have identified altered microbial profiles associated with obesity, insulin resistance, and T2D [10–14]. In agreement with these findings, frequent time zone shifts (jetlag) induce major alterations in overall gut microbiota communities and loss of daytime-dependent oscillation in specific taxa [15]. Importantly, in large human cohorts we showed that microbiota composition and function undergo 24-h rhythmicity and are disrupted in subjects with obesity and/or type 2 diabetes (T2D) [16]. Interestingly, our results in prediabetic patients indicate that arrhythmicity of specific taxa precedes the onset of diabetes and a signature of arrhythmic bacteria predicts T2D risk in populations. Of importance, our recent work on mice identified clocks in cells of the GI tract to be the major regulators of microbial rhythmicity and, therefore, GI homeostasis [17]. Consequently, we hypothesize that intestinal clock-controlled oscillation of the microbiome provides a functional link to metabolic requirements of the host to maintain metabolic health. Here we investigate the impact of circadian disruption on the synchronization of GI clocks and the rhythmicity of microbiota composition and function. Our results show desynchronization of GI clocks in two independent models of circadian disruption, a genetic approach using mice with central circadian dysfunction and an environmental approach using simulated shift work (SSW) on wild type mice. Arrhythmicity of microbial taxa was observed in both models, although microbiota composition differed between experiments. Importantly, arrhythmic bacterial taxa and metabolites identified in both models shared functionalities relevant for metabolic homeostasis of the host. Microbiota transfer further revealed a cross-talk between oscillating taxa and intestinal clocks, highlighting the physiological relevance of microbial rhythms for metabolic health and as therapeutic target.

## 2 Material and methods

### 2.1 Ethics Statement

Experiments were conducted at Technical University of Munich in accordance with Bavarian Animal Care and Use Committee (TVA ROB-55.2Vet-2532.Vet_02-18-14) or were conducted at the University of Lübeck licensed by the Ministry of Agriculture, Environment and Rural Areas (MELUR) of the state of Schleswig-Holstein (project license:42-5/18_Oster).

### 2.2 Mouse models and light conditions

#### 2.2.1 Syt10^cre^-Bmal1^IEC^ ^+/-^ and Syt10^cre^-Bmal1^IECfl/-^ mice

Male SCN-specific *Bmal1* knock-out (Synaptotagmin-10 CRE/wt x *Bmal1fl/-*; referred to as *Bmal1^SCNfl/-^*) mice and their control littermates (Synaptotagmin-10 CRE/wt x *Bmal1+/-*; referred to as *Bmal1^SCN+/-^*) on a genetic C57BL/6J background, mice were generated at the University of Lübeck as described before [18]. Male mice were maintained under a 12 hours light and 12 hours darkness schedule (LD) cycle for 2 weeks (age 8-10 weeks), and sacrificed at the indicated time points during the 2^nd^ day in constant darkness (DD).

#### 2.2.2 Simulated shift work (SSW)

Wild type mice on a genetic C57BL/6J background were bred in house at the Technical University of Munich. Male mice were kept in LD 12:12 cycles (300 lux), with lights turned on at 5am (*Zeitgeber* time (ZT0) to 5pm (ZT12)). Mice were single housed at the age of 8 weeks in running wheel-equipped cages with ad libitum access to chow and water and under specific-pathogen free (SPF) conditions according the FELASA recommendation. To minimize cage-related bias in microbiota composition [19], littermates and litters of comparable age from as few as possible breeding pairs and cages were selected. One set of control males was maintained under a LD cycle for 8 weeks (age 8-16 weeks), whereas another set of male mice was first exposed to for 2 weeks (age 8-10 weeks) of LD and then subjected to SSW conditions for at least 6 weeks. During the experiment mice were exposed to 100lux light intensity and shifted every 5^th^ day by 8 hours. On day 1 of the jet lag, the lights-off time (ZT12) was shifted from 5 pm to 9 am (phase advance paradigm) and from 9 am to 5 pm (phase delay paradigm). Using a short day protocol, we defined day 1 as the first advanced dark period as defined previously [5].

#### 2.2.3 Germ free colonization experiment Transfer experiments

germ-free wild type C57BL6 were gavaged at the age of 10 weeks with cecal microbiota from mixture of cecal content diluted 1:10 in 40% glycerol. Cecal microbiota of 4-5 mice from LD and SSW group were adjusted to 7×10^6^ bacteria/µl and 100µl of were used for gavaging each mouse at ZT13. Germ free recipient mice kept in LD12:12 and were checked weekly for bodyweight changes. After 6 weeks of the gavage, at age 16 weeks, mice were released in constant darkness and sacrificed at the 2^nd^ day at the indicated time point.

### 2.3 Tissue collection

All animals were sacrificed by cervical dislocation followed by decapitation at the age of 16-20 weeks. *Bmal1^SCNfl/-^* mice were sacrificed during the 2^nd^ day of darkness at the indicated circadian time (CT) points. Control mice in the SSW experiment were sacrificed in LD conditions at the indicated *Zeitgeber* time (ZT) and animals undergoing SSW were sacrificed during the 1^st^ day following the final phase advance of SSW at the indicated time point according to the LD control cohort. Tissues were collected and snap frozen using dry ice and stored in −l80 degrees until further processing.

### 2.4 Gut permeability

Gut permeability were measured using Ussing chambers as descibred previously ([20–22]). Briefly, we took 1.5 cm of the proximal colon directly after dissecting the mice. The tissue was cut open and fixed as a flat sheet separating the two halved of the Ussing chamber (six chamber system - Scientific instruments). The tissue was supported from the two sides with carbogen-gassed freshly prepared Krebs buffer (5.4 mM KCl, 114 mM NaCl, 1.2 mM CaCl2,21 mM NaHCO3, 1.2 mM MgCl2, 2.4 mM Na2HPO4, 10 mM glucose, 0.6 mM NaH2PO4, pH 7.4) at 37°C. We added 250ul of 1.7673mM fluorescein to the luminal side, then we determined the fluorescence intensity at 45 and 60 minutes from the buffer on the serosal part, to calculate tissue permeability in cm/s.

### 2.5 Gene expression analysis (qRT-PCR) Quantitative real-time PCR

Snap frozen tissue samples were used to extract RNA samples with Trizol reagent. Next we used 1000ng RNA to synthesize cDNA with cDNA synthesis kit Multiscribe RT (Thermofischer Scientific). We preform qPCR in a Light Cylcer 480 system (Roche Diagnostiscs, Mannheim, Germany) using Universal Probe Library system (UPL) according to manufacturer’s instructions. We used the following primers and probes to measure gene expression: Brain and Muscle ARNT-Like 1 *(Bmal1)* F 5’-ATTCCAGGGGGAACCAGA-’ R 5’-GGCGATGACCCTCTTATCC-3’ Probe 15, Nuclear receptor subfamily 1 group D member 1 *(Rev-erbα)* F 5’-AGGAGCTGGGCCTATTCAC-3’ R 5’-CGGTTCTTCAGCACCAGAG-3’ probe 1, Period 2 *(Per2)* F 5’-TCCGAGTATATCGTGAAGAACG-3’ R 5’-CAGGATCTTCCCAGAAACCA-3’ probe 5, D Site Of Albumin Promoter (Albumin D-Box) Binding Protein (*Dbp*) F 5’-ACAGCAAGCCCAAAGAACC-3’ R 5’-GAGGGCAGAGTTGCCTTG-3’ probe 94, (*Cry1*) F 5’-ATCGTGCGCATTTCACATAC-3’ R 5’-TCCGCCATTGAGTTCTATGAT-3’ probe 85, Glucose transporter 2 (Glut2) F 5’-TTACCGACAGCCCATCCT-3’ R 5’-TGAAAAATGCTGGTTGAATAGTAAAA-3’ probe 3, Fatty Acid Binding Protein 2 (Fabp2) F 5’-ACGGAACGGAGCTCACTG-3’ R 5’-TGGATTAGTTCATTACCAGAAACCT-3’ probe 56, Peroxisome Proliferator Activated Receptor Gamma (Pparg) F 5’-AAGACAACGGACAAATCACCA-3’ R 5’-GGGGGTGATATGTTTGAACTTG-3’ probe 7, Histone Deacetylase 3 (HDAC3) F 5’-GAGAGGTCCCGAGGAGAAC-3’ R 5’-CGCCATCATAGAACTCATTGG-3’ probe 40, Intestinal-type fatty acid-binding protein (Ifabp) 5’-GGTTTCTGGTAATGAACTAATCCAG-3’5’-AAATCTGACATCAGCTTAGCTCTTC-3’ probe 1, the housekeeping gene Elongation factor 1-alpha *(Ef1a)* F 5’-GCCAAT TTCTGGTTGGAATG-3’ R 5’-GGTGACTTTCCATCCCTTGA-3’ probe 67 was used to normalize gene expression.

### 2.6 Nuclear magnetic resonance (NMR)

Body composition (fat, lean mass, free fluid) was measured using a minispec TD-NMR analyser (Bruker Optics, Ettlingen, Germany). Mice were placed in a plastic restrainer and inserted in the minispec for measurements

### 2.7 Energy assimilation

Fecal samples were collected from individual mice over 5 days and dried at 55 °C for another 5 days. Dried fecal pellets were grinded using the TissueLyserII (Qiagen, Retsch, Haan, Germany) and pressed into pellets of 1 gram (technical duplicates). Gross fecal energy content was measured using a 6400 calorimeter (Parr Instrument Company, Moline, IL, USA). Assimilation efficiency was calculated by recording the food intake and feces production over the fecal collection days as indicated in the formula below.

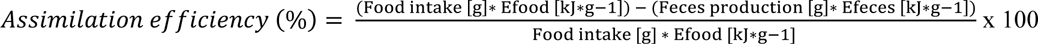

### 2.8 High-Throughput 16S Ribosomal RNA (rRNA) Gene Sequencing and microbial Analysis

Snap-frozen fecal samples was possessed in accordance to slightly mpdified protocol from Godon and colleagues to isolate genomic DNA [23]. DNA was purified with DNA NucleoSpin gDNA columns (Machery-Nagel, No. 740230.250). 24ng DNA was used in a two-step PCR using 341F-ovh and 785r-ov primer to amplify V3-V4 region of 16s rRNA. Sampled were pooled and sequenced in pair-end mode (2×250 bp) on Illumina HISeq using Rapid V2 chemistry, as previously described ({Reitmeier, 2020 #48}). For every 45 samples we included two negative controls of DNA stabilizer without fecal samples to insure reproducibility and control for artifacts. High quality sequence of 16s rRNA with >500 read counts were used for microbial data analysis. FASTQ files were further processed with NGSToolkit (Version 3.5.2_64) with trim score of 5 at both 5’ and 3’ end of R1 and R2 read, then chimera was removed with FASTQ mergepair script of USEARCH. Zero-radius operational taxonomic units (zOTUs) were generated after denoising, deduplicating, clustering and merging quality filtered reads. Here we used zOTUs to have the utmost possible resolution of 16s rRNA sequencing by correcting for sequencing error and identifying sequence with 100% similarity as a unique microbial strain. Taxonomy was assigned based on EZBiocloud database, and RHEA pipeline was used to analyze the data. We aligned the sequence by the maximum likelihood approach wuth MUSCLE from the software MegaX to generate phylogenetic trees and use the online tool Evolview for tree visualization (http://www.evolgenius.info/evolview) [24]. For quantitative analysis, we add spike of 12 artificial DNA that mimics 16s rRNA genes in order to determine 16s rRNA genes copy numbers per gram of fecal sample as previously described [17]

### 2.9 PICRUST 2.0

Metagenomic functionality were predicted using PICRUST2.0. Briefly, based on zOTUs sequence metagenome was constructed to predict functional genes, Normalized zOTU copy numbers were multiplied by the genes for each zOTU. Finally, enzymatic genes were classified to Enzyme Commission (EC) numbers and were assigned to Metacyc pathways. After removing super-classes, and we used Metacyc pathways for LDA effective score (LEFSE) calculation [25] using the online tool (http://huttenhower.sph.harvard.edu/galaxy).

### 2.10 Sample preparation for targeted analysis

Approximately 20 mg of mouse faeces was weighed in a 2 mL bead beater tube (Lysing Matrix D, MP Biomedicals). 1 mL of methanol-based dehydrocholic acid extraction solvent (c=1.3 µmol/L) was added as an internal standard to correct for work-up losses. The samples were extracted 3 times for 20 seconds with 6 m/sec with 30 seconds breaks in using a FastPrep-24 5G bead beating grinder (MP Biomedicals) supplied with a CoolPrep adapter.

### 2.11 Targeted bile acid (BA) measurement

20 µL of isotopically labeled bile acids (ca. 7 µM each) were added to 100 µL of sample extract. Targeted bile acid measurement was performed using a QTRAP 5500 triple quadrupole mass spectrometer (Sciex, Darmstadt, Germany) coupled to an ExionLC AD (Sciex, Darmstadt, Germany) ultrahigh performance liquid chromatography system according to Reiter et al.[26]. Briefly, a multiple reaction monitoring (MRM) method was used for the detection and quantification of the bile acids. An electrospray ion voltage of −4500 V and the following ion source parameters were used: curtain gas (35 psi), temperature (450 °C), gas 1 (55 psi), gas 2 (65 psi), and entrance potential (−10 V). For separation of the analytes a 100 × 2.1 mm, 100 Å, 1.7 μm, Kinetex C18 column (Phenomenex, Aschaffenburg, Germany) was used. Chromatographic separation was performed with a constant flow rate of 0.4 mL/min using a mobile phase consisted of water (eluent A) and acetonitrile/water (95/5, v/v, eluent B), both containing 5 mM ammonium acetate and 0.1% formic acid. The gradient elution started with 25% B for 2 min, increased at 3.5 min to 27% B, in 2 min to 35% B, which was hold until 10 min, increased in 1 min to 43% B, held for 1 min, increased in 2 min to 58% B; held 3 min isocratically at 58% B, then the concentration was increased to 65% at 17.5 min, with another increase to 80% B at 18 min, following an increase at 19 min to 100% B which was hold for 1 min, at 20.5 min the column was equilibrated for 4.5 min at starting. The injection volume for all samples was 1 μL, the column oven temperature was set to 40 °C, and the auto-sampler was kept at 15 °C. Data acquisition and instrumental control were performed with Analyst 1.7 software (Sciex, Darmstadt, Germany).

### 2.12 Targeted short chain fatty acid (SCFA) measurement

The 3-NPH method was used for the quantitation of SCFAs [27; 28]. Briefly, 40 µL of the fecal extract and 15 µL of isotopically labeled standards (ca 50 µM) were mixed with 20 µL 120 mM EDC HCl-6% pyridine-solution and 20 µL of 200 mM 3-NPH HCL solution. After 30 min at 40°C and shaking at 1000 rpm using an Eppendorf Thermomix (Eppendorf, Hamburg, Germany), 900 µL acetonitrile/water (50/50, v/v) was added. After centrifugation at 13000 U/min for 2 min the clear supernatant was used for analysis. The same system as described above was used. The electrospray voltage was set to −4500 V, curtain gas to 35 psi, ion source gas 1 to 55, ion source gas 2 to 65 and the temperature e to 500°C. The MRM-parameters were optimized using commercially available standards for the SCFAs. The chromatographic separation was performed on a 100 × 2.1 mm, 100 Å, 1.7 μm, Kinetex C18 column (Phenomenex, Aschaffenburg, Germany) column with 0.1% formic acid (eluent A) and 0.1% formic acid in acetonitrile (eluent B) as elution solvents. An injection volume of 1 µL and a flow rate of 0.4 mL/min was used. The gradient elution started at 23% B which was held for 3 min, afterward the concentration was increased to 30% B at 4 min, with another increase to 40%B at 6.5 min, at 7 min 100% B was used which was hold for 1 min, at 8.5 min the column was equilibrated at starting conditions. The column oven was set to 40°C and the autosampler to 15°C. Data acquisition and instrumental control were performed with Analyst 1.7 software (Sciex, Darmstadt, Germany).

### 2.13 Statistical analysis

Statistical analysis was perfromed unsing GraphPad Prism, version 9.3.0 (GraphPad Software), R and online platforms (see below). The RHEA pipline (Lagkouvardos) was used to calculate generalized Unifrac distances between sample and consequently to determine microbiota diversity, MDS plots were used to visualize distances between samples [29]. To calculate the cicadian pattern of each 24h period graphs, we used cosine-wave equation: y=baseline+(amplitude·cos(2·π·((x-[phase shift)/24))), with a fixed 24-h period. This equation was used to determine significance of rhythmicity of clock genes, richness, phyla, family and exemplatory profiles of zOTUs. Overall rhythmicity of zOTUs was determined with JTK_CYCLE algorithim [30]. For the manhattan plots JTK_CYCLE was used to calculate amplitude and p-value, and the phase was calculated by cosine-wave regression. Evolview was used for tree visualization (http://www.evolgenius.info/evolview)[24]. To generate heatmaps with the online tool (heatmapper.ca) [31], we sorted the zOTUs or pathways based on the phase of the control group for visualization. The R package SIAMCAT with the function “check.association” [32] was used to generate abundance plots. In order to compare two groups, the non-parametric Mann-Whitney test was used. Two-way ANOVA was used to compare weight gain, clock genes expression in SSW and transfer experiment with Tukey posthoc test for multiple comparison. P-values ≤0.05 were assumed as statistically significant.

## 3 Results

### 3.1 Central clock dysfunction induces circadian desynchronization in the GI tract

Recently we showed that when mice lacking a functional central clock are released into constant darkness (DD), peripheral clocks such as the adrenal, liver, kidney, heart, pancreas, and white adipose tissue gradually desynchronize [6; 9]. As a consequence of system-wide circadian desynchronization, these mice develop obesity and altered glucose metabolism [6]. Of importance, metabolic homeostasis is partially controlled by GI functions regulated by the circadian system [33]. To investigate the degree of circadian desynchronization in peripheral clocks within the GI tract, we compared clock gene expression rhythms in the jejunum, cecum and proximal colon between mice lacking the major clock gene *Bmal1* specifically in the SCN (*Bmal1^SCNfl/-^*) and their littermate controls (*Bmal1^SCN+/-^*) on the 2^nd^ day of DD (**Fig. 1, Table 1**). Circadian rhythmicity analysis revealed that the expression of the core clock genes *Bmal1, Per2* and *Rev-erbα* followed circadian oscillation in the jejunum in both genotypes (cosine-wave regression, control: p=0.004, p=0.02, p=0.03, *Bmal1^SCNfl/-^*: p=0.01, p=0.01, p=0.04) (**Fig. 1A, Table 1**). However, the circadian phases in all clock genes examined in *Bmal1^SCNfl/-^* were significantly advanced (*Bmal1*: 2.7h, *Per2*: 3.6h, *Rev-erbα:* 5.7h). In addition, the baseline of *Rev-erbα* was reduced, *Dbp* did not show significant rhythmicity using cosine regression, but a significant time effect was found in both genotypes by two-way ANOVA analysis (p=0.01). *Cry1* lost rhythmicity in *Bmal1^SCNfl/-^* mice (*Cry1*: p=0.009, p=0.42). In the cecum, all clock genes examined lost rhythmicity in *Bmal1^SCNfl/-^* mice, although a time effect was found for both genotypes by two-way ANOVA (time: *Bmal1* p=0.006, *Per2* p=0.002, *Rev-erbα* p=0.0009, *Dbp* p=0.03, *Cry1* p=0.003) (**Fig. 1B, Table 1)**. In contrast, rhythmicity of *Bmal1, Per2* and *Cry1* gene expression in the proximal colon was undistinguishable between genotypes, and the amplitude of *Rev-erbα* expression was significantly reduced (cosine regression, p=0.02). Similar to results obtained from jejunum, *Dbp* lost rhythmicity in *Bmal1^SCNfl/-^*mice (**Fig. 1C, Table 1**). Altogether, these results suggest that in *Bmal1^SCNfl/-^* mice the jejunal clock free-runs with a reduced amplitude, the cecal clock slowly loses its functionality, whereas the colon clock is functional, albeit with a dampened amplitude. Consequently, these data demonstrate profound disruption of GI clocks in the absence of a functional central clock, which appears at a very early stage following release into constant darkness.

**Figure 1.**
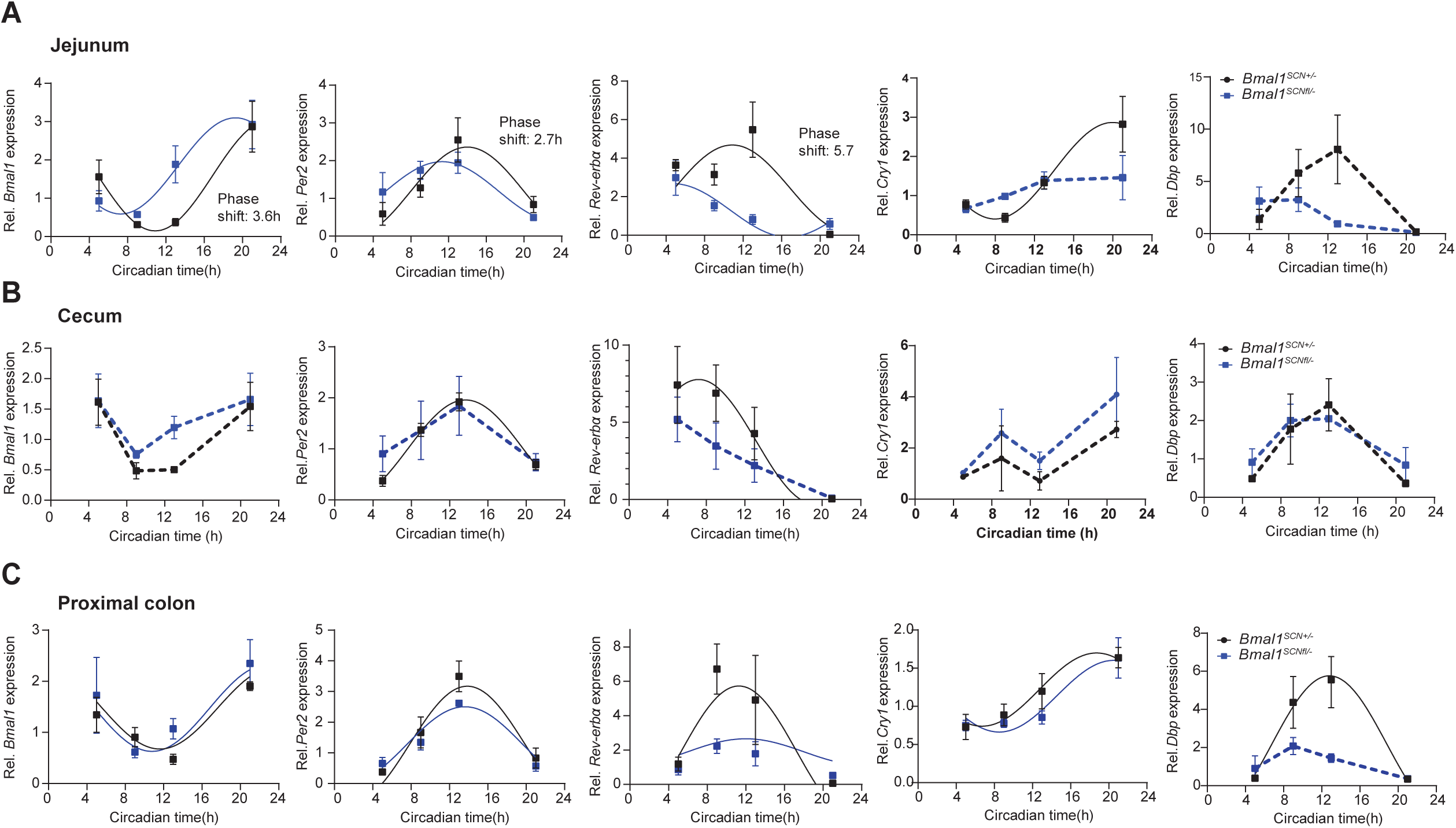
Central clock dysfunction induces circadian desynchronization in the GI tract. Relative expression of core and accessory clock genes in the jejunum (**A**), cecum (**B**), proximal colon (**C**) of *Bmal1^SCNfl/-^* mice (blue) and their controls *Bmal1^SCN+/-^* (black). Significant rhythms according to cosine-wave regression analysis (p-value ≤ 0.05) are visualized with a solid line, while data connected by dashed line indicate arrhythmicity. Significant phase shifts (p ≤ 0.05) are indicated with the number of hours of phase shift. n = 3-4 mice/time point/genotype. Data are represented as mean ± SEM.

**Table 1.** Summary of results for phase, amplitude, baseline and rhythmicity of core and accessory clock gene expression based on cosine regression analysis in the GI tract of Bmal1^SCNfl/-^ mice and control. Bold p-values indicate significant difference between genotype.

| Gastrointestinal tissue | Gene | Group | Rhythmicity<br>(P. value) | Phase<br>shift<br>(P. value) | Amplitude<br>difference<br>(P. value) | Baseline<br>difference<br>(P. value) |
| --- | --- | --- | --- | --- | --- | --- |
| Proximal colon | *Bmal* | *Bmall*<sup>SCN</sup><br>+/- | 0.01 | 0.53 | 0.67 | 0.22 |
|  |  | *Bmall*<sup>SCN</sup><br>fl/- | 0.04 |  |  |  |
|  | *Per2* | *Bmall*<sup>SCN</sup><br>+/- | 0.004 | 0.56 | 0.13 | 0.25 |
|  |  | *Bmall*<sup>SCN</sup><br>fl/- | 0.001 |  |  |  |
|  | ***Rev-erba*** | *Bmall*<sup>SCN</sup><br>+/- | 0.03 | 0.98 | **0.02** | 0.07 |
|  |  | *Bmall*<sup>SCN</sup><br>fl/- | 0.03 |  |  |  |
|  | *Cry1* | *Bmall*<sup>SCN</sup><br>+/- | 0.01 | 0.27 | 0.94 | 0.47 |
|  |  | *Bmall*<sup>SCN</sup><br>fl/- | 0.01 |  |  |  |
|  | ***Dbp*** | *Bmall*<sup>SCN</sup><br>+/- | **0.008** | 0.39 | **0.002** | **0.04** |
|  |  | *Bmall*<sup>SCN</sup><br>fl/- | **0.07** |  |  |  |
| Cecum | *Bmal* | *Bmall*<sup>SCN</sup><br>+/- | 0.06 | 0.53 | 0.34 | 0.3 |
|  |  | *Bmall*<sup>SCN</sup><br>fl/- | 0.33 |  |  |  |
|  | ***Per2*** | *Bmall*<sup>SCN</sup><br>+/- | **0.00006** | 0.54 | 0.37 | 0.59 |
|  |  | *Bmall*<sup>SCN</sup><br>fl/- | **0.31** |  |  |  |
|  | ***Rev-erba*** | *Bmall*<sup>SCN</sup><br>+/- | **0.02** | 0.59 | 0.29 | 0.12 |
|  |  | *Bmall*<sup>SCN</sup><br>fl/- | **0.08** |  |  |  |
|  | *Cry1* | *Bmall*<sup>SCN</sup><br>+/- | 0.37 | 0.76 | 0.74 | 0.15 |
|  |  | *Bmall*<sup>SCN</sup><br>fl/- | 0.52 |  |  |  |
|  | ***Dbp*** | *Bmall*<sup>SCN</sup><br>+/- | 0.056 | 0.78 | 0.34 | 0.45 |
|  |  | *Bmall*<sup>SCN</sup><br>fl/- | 0.1 |  |  |  |
| Jejunum | *Bmal* | *Bmall*<sup>SCN</sup><br>+/- | 0.004 | 0.02 | 0.64 | 0.42 |
|  |  | *Bmall*<sup>SCN</sup><br>fl/- | 0.01 | Phase<br>shift=2.7 |  |  |
|  | *Per2* | *Bmall*<sup>SCN</sup><br>+/- | 0.02 | 0.04 | 0.28 | 0.83 |
|  |  | *Bmall*<sup>SCN</sup><br>fl/- | 0.01 | Phase<br>shift=3.6 |  |  |
|  | *Rev-erba* | *Bmall*<sup>SCN</sup><br>+/- | 0.03 | 0.03 | 0.28 | 0.04 |
|  |  | *Bmall*<sup>SCN</sup><br>fl/- | 0.04 | Phase<br>shift=5.7 |  |  |
|  | *Cry1* | *Bmall*<sup>SCN</sup><br>+/- | 0.009 | 0.33 | 0.07 | 0.11 |
|  |  | *Bmall*<sup>SCN</sup><br>fl/- | 0.42 |  |  |  |
|  | *Dbp* | *Bmall*<sup>SCN</sup><br>+/- | 0.056 | 0.08 | 0.08 | 0.14 |
|  |  | *Bmall*<sup>SCN</sup><br>fl/- | 0.06 |  |  |  |

### 3.2 Disruption of microbiota rhythmicity in SCN-specific *Bmal1*-deficient mice

GI clocks are dominant regulators of circadian microbiome fluctuations and thereby balance GI homeostasis, as previously shown by us [17]. This prompted us to determine whether circadian desynchronization in GI tissues in *Bmal1^SCNfl/-^* mice affects circadian microbiota composition and function. Indeed, 16s rRNA analysis of fecal samples revealed significant clustering according to genotype (**Fig. 2A**), suggesting a different microbiota composition in *Bmal1^SCNfl/-^*mice. Moreover, circadian rhythmicity in community diversity (species richness) observed in control mice was abolished in *Bmal1^SCNfl/-^*mice, although Generalized UniFrac distance (GUniFrac) quantification to CT1 identified a time difference in both genotypes (two-way ANOVA, p=0.0037) (**Fig. 2B**). Relative abundance of the two major phyla, *Firmicutes* and *Bacteroidetes*, showed circadian rhythmicity with similar patterns in both genotypes (**Fig. 2C**). However, previous research, including from our own group, showed that rhythmicity in relative abundance can be masked due to oscillations of highly abundant taxa [17; 34]. Thus, we used synthetic DNA spikes to determine quantitative microbiota composition as previously described [35]. Indeed, both phyla lost rhythmicity in quantitative abundance in *Bmal1^SCNfl/-^*mice compared to controls (**Fig. 2C**). Central clock disruption led to loss of rhythmicity of the families *Lactobacillaceae* and *Clostridiales* independent of the analysis (**Suppl. Fig.1A**). Then we set out to determine rhythmicity of zero-radius OTUs (zOTUs) after removal of low-abundance taxa (mean relative abundance < 0.1%; prevalence < 10%). The heatmaps illustrate disrupted circadian oscillations of zOTUs in *Bmal1^SCNfl/-^* mice for both analyses (**Fig. 2D, Suppl. Fig. 1B)**. The amount of rhythmic zOTUs was reduced by three quarters in mice with SCN-specific *Bmal1* deficiency (JTK_CYCLE, adj. p-value < 0.05) (**Fig. 2E, Suppl. Fig.1C, Suppl. Table. 1)**. For example, we identified zOTUs which lost rhythmicity in *Bmal1^SCNfl/-^* mice predominantly belonging to mucus foragers (*Muribaculaceae*) and to the secondary bile acid and SCFA producing family *Ruminococcaceae* [36; 37] (**Fig. 2F, Suppl. Fig.1D**). In particular, SCFA producing taxa, including *Faecalibaculum* and *Agathobaculum* [38], were arrhythmic in *Bmal1^SCNfl/-^* mice (**Fig. 2F, G, Suppl. Fig.1D, E**). Of note, bacteria belonging to *Alloprevotella, Muribaculaceae* and *Faecalibaculum* lost rhythmicity and additionally differed in their abundance between genotypes (**Suppl. Fig. 1F**).

**Figure 2.**
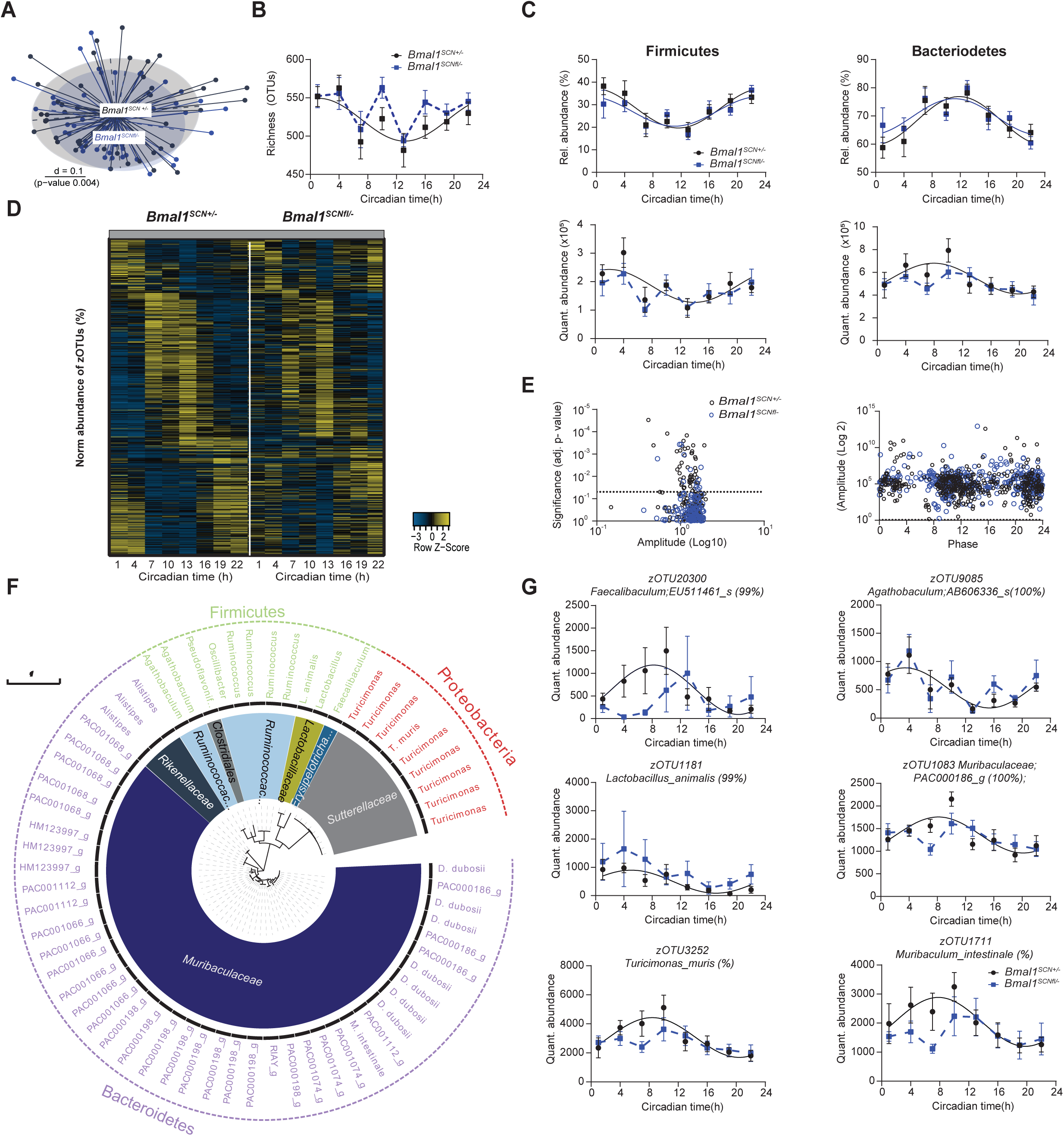
Disruption of microbiota rhythmicity in SCN-specific *Bmal1*-deficient mice. (**A**) Beta-diversity MDS plot based on generalized UniFrac distances (GUniFrac) of fecal microbiota stratified by genotype. (**B-C**) Circadian profile of alpha diversity (**B**) and the relative and absolute abundance of major phyla (**C**). (**D**) Heatmap illustrating the relative abundance of 412 zOTUs (mean relative abundance > 0.1%; prevalence > 10%). Data are ordered based on the zOTUs phase in the controls and normalized based in the peak of each zOTU. (**E**) Significance and amplitude (based on JTK_CYCLE) of all zOTUs (left) and phase (based on cosine regression) distribution (right) in both genotype. Dashed line represent adj. p-value = 0.05 (JTK_CYCLE). (**F**) Taxonomic tree of zOTUs losing rhythmicity in *Bmal1^SCNfl/-^* mice based on quantitative analyses. Taxonomic ranks were indicated as phylum (outer dashed ring), families (inner circle) and genera (middle names). Each zOTU is represented by individual branches. (**G**) Circadian profiles of absolute abundance of example zOTUs losing rhythmicity in *Bmal1^SCNfl/-^* mice. Significant rhythms according to cosine-wave regression analysis (p-value ≤ 0.05) are visualized with a solid line, while data connected by dashed line indicate arrhythmicity. n = 6 mice/time point/genotype. Data are represented as mean ± SEM.

### 3.3 SCN clock-controlled microbial functions balance metabolic homeostasis

To address the potential physiological relevance of microbial rhythmicity we performed PICRUST 2.0 analysis on zOTUs which lost rhythmicity in *Bmal1^SCNfl/-^* mice [39]. SCN clock-deficient mice develop adiposity and impaired glucose handling [6]. In this context, genotype differences and loss of rhythmicity was observed in predicted pathways related to sugar metabolism, SCFA fermentation and fatty acid metabolism (**Fig. 3A, Suppl. Fig. 2A**). Targeted metabolite analysis further revealed that alterations in taxa identified in *Bmal1^SCNfl/-^*mice led to changes in key bacterial products involved in sugar and lipid signaling, such as SCFAs and (BAs (**Fig. 3B-F, Suppl. Fig. 2B-D)**. In particular, propionic acid, important for lipid metabolism [40], showed reduced levels in *Bmal1^SCNfl/-^* mice (**Fig. 3B)**. Moreover, branched-chain fatty acids including isovaleric acid, isobutyric acid and 2-methylbutyric acid were reduced in *Bmal1^SCNfl/-^* mice, whereas total SCFA concentrations were undistinguishable between genotypes (**Fig. 3B, Suppl. Fig. 2B)**. Rhythmicity of total SCFAs as well as of major microbial derived products such as acetic acid, propionic acid and lactic acid was absent in *Bmal1^SCNfl/-^* mice (cosine regression, control: p=0.003, p=0.001, p=0.02, p=0.0009, *Bmal1^SCNfl/-^*: p=0.32, p=0.5, p=0.49, p= 0.93) (**Fig. 3C**). Of note, other SCFAs, including butyric acid and valeric acid, showed rhythmicity in both genotypes (cosine regression, control p=0.0001, p=0.007, *Bmal1^SCNfl/-^*, p=0.02, p=0.01, respectively) (**Suppl. Fig. 2C**). In addition, BA concentrations were altered in mice lacking a functional central clock (**Fig. 3D**, **Suppl. Fig. 2D**). For example, 6-ketolithocholic acid concentrations were reduced, whereas concentrations of b-muricholic acid and tauro-a-muricholic acid were significantly elevated in *Bmal1^SCNfl/-^* mice (**Fig. 3D**). Although other BAs measured had comparable concentrations in both genotypes, rhythmicity of various BAs was disrupted in *Bmal1^SCNfl/-^* including, 7-sulfocholic acid, ursodeoxycholic acid, taurocholic acid and allolithocholic acid (**Fig. 3E, F, Suppl. Fig. 2D**), suggesting altered fat and cholesterol metabolism [41].

**Figure 3.**
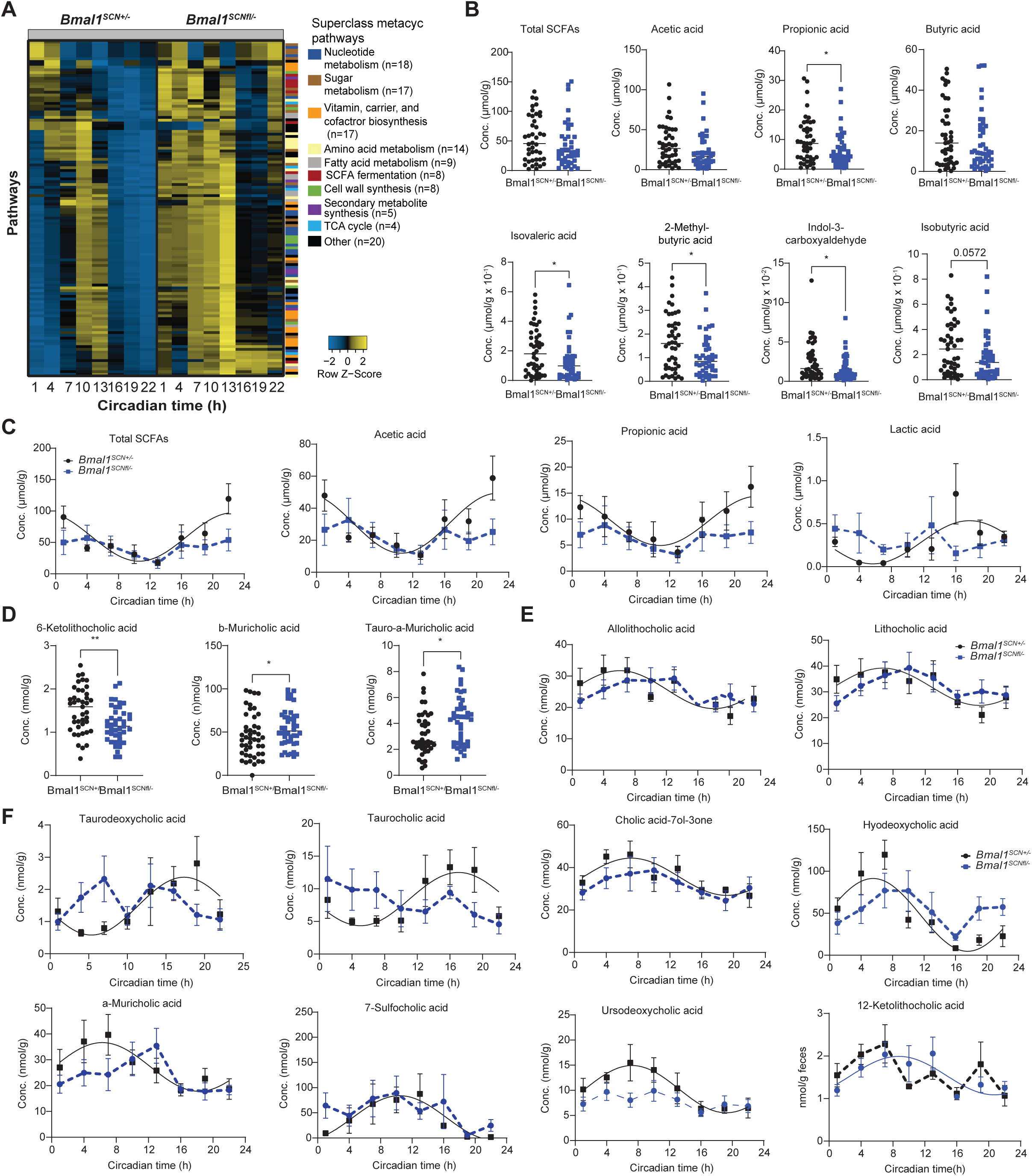
SCN clock-controlled microbial functions balance metabolic homeostasis. (**A**) Heatmap representing MetaCyc Pathways predicted by PICRUST2.0 from zOTUs losing rhythmicity in *Bmal1^SCNfl/-^*mice. Pathways are ordered by the phase of the control and normalized to the peak abundance of each pathway. We color-coded the pathways according to their sub-classes. (**B**) Fecal SCFA concentrations in both genotype. (**C**) Circadian profiles of fecal SCFA. (**D**) Fecal bile acid concentrations in both genotype. (**E-F**) Circadian profiles of fecal bile acids. Significant rhythms according to cosine-wave regression analysis (p-value ≤ 0.05) are visualized with a solid line, while data connected by dashed line indicate arrhythmicity. Mann Whitney U test was used to assess concentration difference. n = 6 mice/time point/genotype. Data are represented as mean ± SEM. Significance * p ≤ 0.05, ** p ≤ 0.01, *** p ≤ 0.001, **** p ≤ 0.0001

Taken together, our results highlight the importance of the central clock in synchronizing peripheral clocks located in GI tissues. In addition, these results show for the first time loss of microbial taxa and their functional outputs, in particular SCFAs and BAs, in mice lacking central lock function, which associates with adiposity and impaired glucose metabolism in these animals [6].

### 3.4 Simulated shift work induces circadian desynchrony between GI clocks

Epidemiological and experimental studies indicate that frequent circadian desynchronization increases the risk of developing metabolic diseases and weight gain [42; 43], similar to the phenotype observed in central clock-deficient mice [6]. Circadian desynchronization among tissue clocks, as observed in mice lacking the central clock [9], can be induced by misalignment between internal and environmental time, such as during jetlag or shift work [5]. To investigate whether shift work induces circadian desynchrony among GI clocks similar to the effects of a loss of central clock function in *Bmal1^SCNfl^*^/-^ mice, wild type mice were exposed to phase shifts of 8 hours every 5^th^ day for 6-8 weeks to SSW (**Fig. 4A**). The activity profiles gradually advanced during the first days in SSW (**Fig. 4A, B**). In particular, in comparison to the LD profiles before SSW and the control cohort kept in LD, the activity onset advanced by less than 3 hours at the 1^st^ day (**Fig. 4A, B)**. This resulted in an equal distribution of activity between prior day and night, although total activity was unaffected (**Fig. 4B, Suppl. Fig. 3A**). In line with previous studies, mice in SSW significantly increased their body weight (P <0.0001) [15] **(Fig. 4C)**. In addition, colon permeability was enhanced at CT13 during SSW, although no difference in energy assimilation or total food intake was detected (**Fig. 4D-F**).

**Figure 4.**
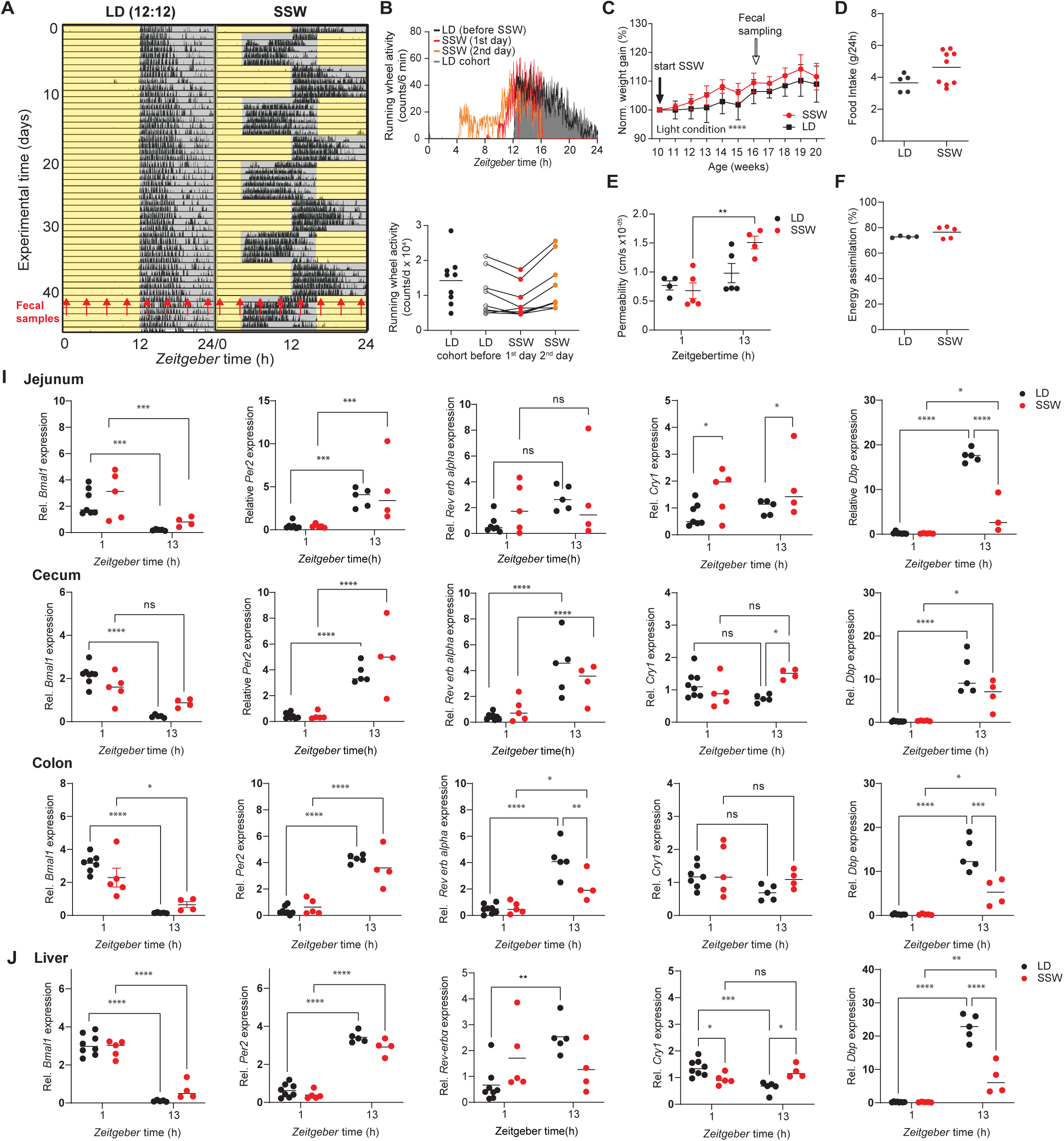
Simulated shift work induces circadian desynchrony between GI clocks. (**A**) Representative actogram of a control mouse in 12-hour light/12-hour dark (LD) and under simulated shift work (SSW) condition. Tick marks represent running wheel activity. Yellow and grey shadings represent light and darkness respectively. Red arrows indicate fecal sample collection time points. (**B**) Diurnal total wheel-running activity profiles (top) and 24-h summary (bottom). (**C**) Normalized body weight gain of mice in SSW and LD condition. Total daily food intake (**D**), gut permeability (**E**) and energy assimilation (**F**). (**I-J**) Relative expression of core and accessory clock genes in GI tract (**I**) and liver (**J**) of WT mice in SSW (red) and their LD controls (black). N = 4-5 mice/time point/light condition. Data are represented as mean ± SEM. Mann Whitney U test was used to assess food intake and energy assimilation differences. Two-way ANOVA was used to assess the change in body weight and gene expression. Significance * p ≤ 0.05, ** p ≤ 0.01, *** p ≤ 0.001, **** p ≤ 0.0001

Differences in the resetting speed of circadian clocks and between clock genes within the same tissue have been reported [5]. To test whether GI clocks are affected differentially by SSW, clock gene expression in GI tissues was measured at ZT1 and ZT13 (1 and 13 hours after the lights on in controls). Indeed, diurnal expression of clock genes in GI tissues and the liver as control was differentially affected at the 1^st^ day during the last phase advance in SSW (**Fig. 4A, I, J**). Although *Bmal1* and *Per2* in the liver, jejunum and proximal colon showed daytime dependent expression in both genotypes, *Dbp*, *Cry1* and *Rev-erbα* were affected only in specific tissues. For example, *Dbp* was dramatically reduced at ZT13 in the liver and jejunum of mice exposed to SSW, whereas no daytime effect, but enhanced expression during SSW, was found for *Dbp* (**Fig. 4I, J**). In contrast, in the cecum daytime differences of *Bmal1* were absent and *Cry1* significantly enhanced its expression at ZT13 during SSW, while *Per2*, *Reverbα* and *Dbp* were unaffected (**Fig. 4I**). Moreover, in the colon of mice undergoing SSW, a time difference in the expression of almost all clock genes examined (except of *Cry1*) was found, although *Bmal1*, *Rev-Erbα* and *Dbp* expression was significantly suppressed at ZT13 (**Fig. 4I**). These results indicate that all peripheral clocks examined were in different resetting stages of the phase advance and consequently circadian desynchronization was evident between GI clocks.

### 3.5 Simulated shift work disrupts rhythmicity of microbiota composition and function

Previous research, including from our own group, indicates that changes in environmental conditions can modify microbial community composition and cause arrhythmicity of specific taxa [15; 17; 44]. In accordance, we found significantly different fecal microbial communities between mice exposed to LD and SSW conditions (p=0.014) (**Fig. 5A**). Rhythmicity of GUniFrac distance quantification as well as the relative and quantitative abundance of major phyla and families was phase shifted in line with the advanced behavioral rhythm (**Fig. 4A, B**, **Fig. 5B-D, Suppl. Fig. 3B, C**). Importantly, the quantitative abundance of *Bacteroidetes* lost rhythmicity in SSW **(Fig. 5C)**. Heatmaps of bacterial abundances over the course of the 24-hour day illustrate phase advanced rhythms of zOTUs during SSW independent of the analysis (**Suppl. Fig. 3D, E**). Moreover, arrhythmicity was identified during SSW in ∼50% of all rhythmic zOTUs in LD conditions, including *Lactobacillus, Ruminococcus* and *Odoribacter* (**Fig. 5E, G, Suppl. Fig. 3D-F, Suppl. Table 1**). zOTUs which lost rhythmicity in quantitative and relative analyses included taxa belonging to *Eubacterium, Bacteroides* and *Ruminococcus* (**Fig. 5G, H**, **Suppl. Fig. 3G, Suppl. Table 1**). The phase of the remaining rhythmic zOTUs in SSW advanced by 3.7 - 6.4h, including the genera *Alistipes, Duncaniella, Roseburia, Oscillibacter* and the family *Lachnospiraceae,* (**Fig. 5F, Suppl. Fig. 3D, F, Suppl. Table 1**). Of note, the average abundance of arrhythmic zOTUs belonging to the *Ruminococcaceae* and *Muribaculaceae* families as well as the genus *Lactobacillus* significantly differed between SSW and LD conditions (**Fig. 5G, Suppl. Fig 3F**) in accordance with results obtained from mice exposed to chronic jetlag or sleep deprivation [15; 44-46].

**Figure 5.**
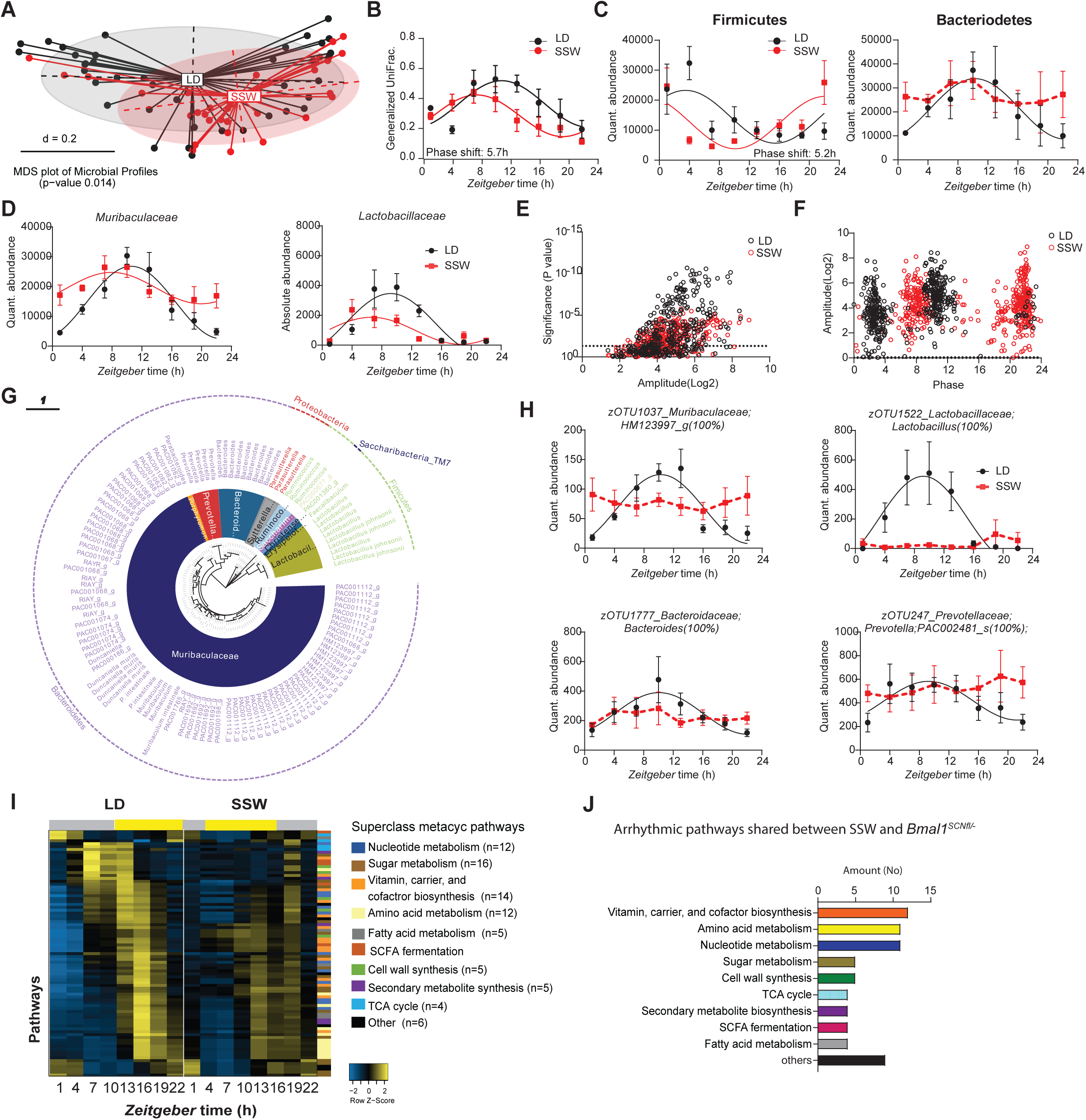
Simulated shift work disrupts rhythmicity of microbiota composition and function. (**A**) Beta-diversity MDS plot based on generalized UniFrac distances (GUniFrac) of fecal microbiota stratified by light condition. (**B**) Circadian profile of generalized unifrac distance normalized towards ZT1 of the controls. (**C-D**) Circadian profiles of the absolute abundance of major phyla (**C**) and families (**D**). (**E**) Significance and amplitude (based on JTK_CYCLE) of all zOTUs (**E**) and phase (based on cosine regression) distribution (**F**) in both genotype, dashed line represents adj. p-value = 0.05 (JTK_CYCLE). (**G**) Taxonomic tree of zOTUs losing rhythmicity in SSW based on quantitative analyses. Taxonomic ranks were indicated as phylum (outer dashed ring), then family (inner circle) and genera (middle names). Each zOTU is represented by individual branches. (**H**) Circadian profiles of absolute abundance of example zOTUs losing rhythmicity in SSW. (**I**) Heatmap representing MetaCyc Pathways predicted by PICRUST2.0 from zOTUs losing rhythmicity in SSW. Pathways are ordered by the phase of the control and normalized to the peak abundance of each pathway. We colored the pathways according to their sub-classes. (**J**) Bar chart representing the number of shared pathways losing rhythmicity in SSW and *Bmal1^SCNfl/-^* mice. Significant rhythms according to cosine-wave regression analysis (p-value ≤ 0.05) are visualized with a solid line, while data connected by dashed line indicate arrhythmicity. Significant phase shifts (p ≤ 0.05) are indicated with the number of hours of phase shift. n = 4-5 mice/time point/genotype. Data are represented as mean ± SEM.

To evaluate whether GI clock desynchronization during SSW might have induced similar disturbance of microbial oscillations as observed in mice with central clock disruption, we analyzed rhythmicity of the microbiome in mice undergoing SSW. Of note, overall microbiota composition was not comparable between these two experiments performed in different animal facilities (**Suppl. Fig. 3H**). However, this is in accordance with frequent reports illustrating that the housing situation dramatically influences microbiota composition [47]. To consider microbiota function rather than composition, we performed PICRUST analysis of zOTUs which lost rhythmicity in SSW **(Fig. 5H**). Their predicted functionality was then compared to results obtained from arrhythmic taxa identified in *Bmal1^SCNfl/-^* mice (**Fig. 3A, Fig. 5H, Suppl. Fig. 3I)**. Independent of the approach of inducing circadian desynchronization, disrupted rhythmicity and changes in abundance were found in pathways related to amino acids, fatty acids as well as sugar metabolism and SCFA fermentation **(Fig. 5I)**, suggesting a functional link between circadian microbiota regulation and GI physiology.

### 3.6 Simulated shift work-associated microbiota promote weight gain and suppress GI clocks

In order to directly investigate the effect of SSW-induced arrhythmicity of the microbiome on the host, we performed cecal microbiota transfer from donor mice undergoing 6 weeks of SSW and controls kept in LD into germ-free (GF) wild type recipients (**Fig. 6A**). Mice receiving SSW-associated microbiota significantly increased their body weight (**Fig. 6B**), in line with observations following fecal microbiota transplantation from mice exposed to chronic jetlag [15]. Interestingly, 6 weeks after transfer, body weight as well as most organ weights were undistinguishable between recipients (**Fig. 6A-C)**, indicating that microbial alterations are temporary in rhythmic hosts. Of note, an increased cecum weight was observed even 6 weeks after transfer (**Fig. 6C**). Microbial derived products, especially SCFAs and BAs have been described to alter clock gene expression in GI tissues [48; 49]. This prompted us to measure clock gene expression in recipients as well as in GF controls. Indeed, mice receiving SSW- associated microbiota showed altered GI clock gene expression 6 weeks after the transfer (**Fig. 6D**). Although most clock genes examined in the proximal colon fluctuated between daytimes independent of the genotype of the donor, *Per2* expression was highly suppressed at ZT13 and the daytime difference of *Rev-erbα* expression in controls was absent in mice receiving SSW- associated microbiota (**Fig. 6D**). Similarly, *Per2, Cry1* and *Dbp* expression in jejunum as well as *Per2*, *Rev-erbα* and *Dbp* expression in cecum was suppressed at ZT13 in mice receiving SSW-associated microbiota. Dampened daytime differences in GI clock gene expression followed similar trends than observations made in donor mice exposed to SSW and in GF mice (**Fig. 4I, Fig. 6D**). These results suggest that the microbiome can at least partly transfer the GI clock phenotype from the donor to the host and thus directly impact GI physiology. In mice receiving SSW-associated microbiota, we then investigated the effect of clock gene suppression on clock-controlled genes related to glucose and fat metabolism, such as *Fabp2*, *Hdac3, Ifab, Glut2* and *Ppary* [50–52]. Indeed, in the jejunum, suppressed expression was found for *Fabp2* involved in lipid uptake [52] and *Glut2* a regulator for glucose uptake [53]. In the colon, enhanced expression was found for *Ppary,* a transcriptional regulator of glucose and lipid metabolism [50] (**Fig. 6E, F**) and SCFAs were shown to modulate the metabolic state of the host through PPARs [54]. Altogether, these results demonstrate the physiological relevance of the GI clock-microbiome crosstalk, specifically for maintenance of the host’s metabolic health.

**Figure 6.**
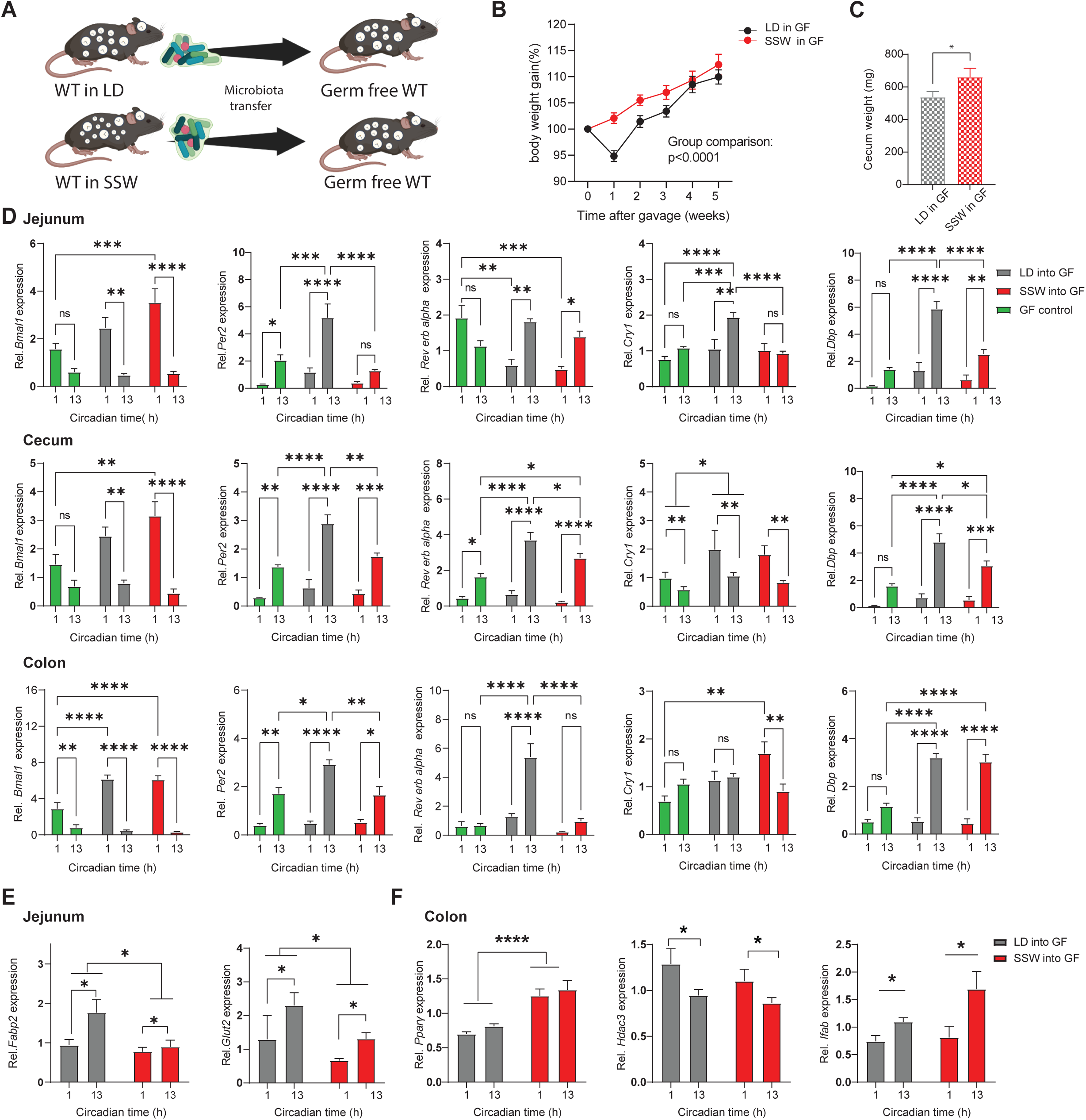
Simulated shift work-associated microbiota promote weight gain and suppress GI clocks. (**A**) Schematic illustration of cecal microbiota transfer from SSW and LD donors (n=4-5) intro germ free wild type mice. (**B**) Normalized body weight gain of recipient mice. (**C**) bar chart illustrated cecum weight in recipient mice. (**D-E**) Relative expression of clock genes (**D**) and clock controlled gene (**E, F**) in the GI tract of germ free mice (green), germ free receiving SSW (red) and LD controls (black) microbiota. N = 5-6 mice/time point/light condition. Data are represented as mean ± SEM. Mann Whitney U test was used to assess the different in cecum weight. Two-way ANOVA was used to assess the change in body weight and differences in gene expression. Significance * p ≤ 0.05, ** p ≤ 0.01, *** p ≤ 0.001, **** p ≤ 0.0001

## 4 Discussion

Mice with central clock dysfunction were shown to develop a metabolic phenotype and desynchrony in peripheral clocks, such as the adrenal, the liver, the heart, the pancreas and eWAT [6; 9]. In addition, we provide evidence that GI clocks desynchronize in the absence of a functional central clock. Moreover, we demonstrate that desynchronization among GI clocks also appears in wild type mice exposed to SSW conditions. Our results comply with alterations in colonic clock gene expression following chronic jet lag [15] and suggest that GI clock desynchrony is a common feature during circadian disruption. Of note, distinct sections of the GI circadian system responded differentially to circadian disturbances, which was evident in the genetic model and during environmentally induced circadian disruption. Considering previous research indicating a temporal phase gradient of clock gene rhythms along the gut cranio-caudal axis [55], the circadian response to circadian disturbance might differ between gut sections. However, 24-hour profiling of clock gene expression over multiple days would be necessary to compare the kinetics of resetting between intestinal tissue clocks.

Recently, we identified that GI clocks are prominent drivers of gut microbiota rhythmicity [17]. Consequently, arrhythmicity of the microbiota observed in mice with central clock disruption and in mice kept in SSW was likely induced by desynchronization among GI clocks. Indeed, in line with recent results obtained from mice with dysfunctional intestinal clocks [17], taxa belonging to the families *Rikenellaceae, Ruminococcaceae* and *Muribaculaceae* as well as to the genera *Lactobacillus* and *Alistipes* lost rhythmicity in *Bmal1^SCNfl/-^* mice and in mice undergoing SSW. Of note, disruption of rhythmicity was more severe in mice lacking a functional central clock. Here, arrhythmicity was found in microbial diversity and on the level of phyla and families. During SSW the abundance of the phylum *Bacteroidetes* and, thus, a substantial amount of taxa remained rhythmic, although with an advanced phase. This discrepancy between both models might be explained by the arrhythmic food intake behaviour documented in *Bmal1^SCNfl/-^* mice upon release in DD [18], whereas in SSW the daily patterns of food intake were rhythmic but phase shifted. Manipulating the timing of food intake has been shown to phase shift specific taxa belonging to *Alistipes, Lactobacillus* and *Bacteroides* [15; 17]. Therefore, a phase-advanced food intake rhythm in SSW could have changed the phase of bacterial oscillations. Nevertheless, a substantial amount of taxa lost rhythmicity upon exposure to SSW and were also found to lose rhythmicity in mice with SCN-specific and GI clock disruption [17], indicating that loss of synchrony between GI clocks may be responsible for microbial arrhythmicity during circadian disruption.

Recently we discovered a link between microbiota rhythmicity, obesity and T2D development in humans [16], suggesting that microbial rhythms may play a causative role for disease development. Accordingly, transfer of microbiota from an obese human donor as well as from lean donors undergoing jetlag induces an obesity-associated phenotype in GF recipient mice [15; 45; 56]. However, these studies did not address whether obesity associated loss of microbial rhythmicity or general changes in abundance of bacteria are the underlying cause. Transfer experiments using mouse models with circadian dysfunction provide direct evidence for the physiological relevance of microbiota rhythms for metabolic health. For example, transfer of arrhythmic microbiota from gut-clock deficient mice disrupts GI homeostasis in recipient animals [17], and microbiota from mice exposed to environmentally induces circadian disruption promoting body weight gain in wild type mice. Similar results were obtained following microbiota transfer from jet lagged mice [15]. Together, these results suggest that on top of peripheral clock disruption in the fat and liver[6; 9], the rhythmicity of the microbiome is a critical factor for the development of metabolic disease.

GI metabolism is strongly influenced by bacterially derived products, such as SCFAs and BAs [57; 58]. After both genetic and environmental circadian disruption, loss of microbial rhythmicity was reflected by arrhythmicity of predicted microbial functionality, such as SCFA fermentation, as well as sugar, fatty acid and amino acid metabolism. Targeted metabolite analysis further confirmed lack of rhythmicity of key microbial derived products in *Bmal1^SCNfl/-^* mice, namely SCFAs and BAs. For example, arrhythmicity was found for the SCFAs Propionic acid and Acetic acid. Both play a major role in fat and glucose metabolism and are capable in preventing diet induced obesity and insulin resistance [40]. Additionally, alterations in either rhythmicity or abundance of taurine-conjugated bile acids as well as the secondary BA Ursodeoxycholic acid were observed. These metabolites are known to impact signaling through the nuclear bile acid receptor FXR, resulting in the transcription of target genes important for lipid and glucose homeostasis (reviewed by [41]). Importantly, bacterial metabolites, such as SCFAs and BAs, are controlled by the circadian clock, and alterations in SCFA and BA oscillations were previously reported in mice exposed to chronic jet lag and in GI clock deficient animals [17; 59]. Loss of rhythmicity of SCFAs as well as BAs which are both involved in sugar and fatty acid metabolism (reviewed by [60]) might alter metabolic functionalities of the host following circadian disruption, since both bacterial products are known to balance host metabolism (reviewed by [61]). In this regard, we previously reported an increased body weight gain, when *Bmal1^SCNfl^*^/-^ mice were kept in DD for multiple weeks [6]. Of importance, loss of microbiota rhythms and subsequent microbial functions predominantly involved in glucose and lipid metabolism, such as Ursodeoxycholic acid, Propionic acid and Acetic acid [57; 62; 63], were already found at the 2^nd^ day of DD and thus precede the obesity phenotype reported in these mice. Consequently, the observed microbial changes might represent an early event in the development of the metabolic phenotype of *Bmal1^SCNfl^*^/-^ mice [6].

Interestingly, shift work associated bacteria directly affect the host’s GI clock function. In particular, GI clock dysregulation in donor mice following circadian disruption was partly reflected in recipients. For example, suppression of daytime differences in colonic *Rev-Erbα* and *Dbp* expression in jejunum was evident in both donor and recipient, indicating that microbiota transfer the circadian phenotype from the donor to recipients. Peripheral circadian clocks are known to control organ functions through regulation of tissue-specific CCGs ([4]). Accordingly, GI clock disruption in recipients altered the expression levels of CCGs in jejunum and colon, such as *Fabp2, Glut2* and *Ppary*, both involved in glucose and fat metabolism [50; 53]. The mechanisms linking microbiota rhythms with functions of the GI tissue likely involve local epithelial-microbial interactions. Indeed, SCFAs and BAs have been reported to directly impact rhythmicity in intestinal epithelial cells and affect metabolic responses of the host [48; 59; 64; 65]. Consequently, arrhythmicity of the transferred microbiota likely resulted in arrhythmicity of bacterial products, capable to alter GI clock function and, subsequently, metabolic CCGs. Therefore, our results provide first mechanistic insights into microbiota-dependent metabolic abnormalities during circadian disruption.

## 5 Conclusions

Taken together, the comparison of two models of genetic and environmentally induced circadian disruption revealed shared disruption at the level of GI clocks and identified microbial taxa and their functionalities involved in metabolic abnormalities of the host. Further, microbial alterations during SSW appear to be causal for the metabolic phenotype of the host. Our data provide first evidence that molecular alterations of GI clock function during circadian disruption are transferrable between organisms through the microbiome. Thereby our data highlight the intestinal clock-bacteria dialogue as a potent underlying factor in the development of metabolic diseases in humans exposed to circadian disruption due to their lifestyle.

## 6 Author contribution

SK conceived and coordinated the project. BA, VP, MH, YN and EG performed mouse experiments and fecal samples collection. YN and MH measured epithelial membrane properties. MH conducted bomb calorimetry and NMR. SK and MH analyzed activity and food intake behavior. BA and MH performed 16S rRNA gene sequencing and bioinformatics analysis. BA analyzed gene expression, predicted microbial functionality and conducted germ free mouse colonization. KK, MG and BA performed targeted metabolomics and data analyses. SK supervised the work and data analysis. SK, HO and DH secured funding. BA, SK and MH wrote the manuscript. All authors reviewed and revised the manuscript.

## 7 Funding

SK was supported by the German Research Foundation (DFG, project KI 19581) and the European Crohńs and Colitis Organisation (ECCO, grant 5280024). SK and DH received funding by the Funded by the Deutsche Forschungsgemeinschaft (DFG, German Research Foundation) – Projektnummer 395357507 – SFB 1371). HO was funded by the DFG (project OS353-11/1).

## 8 Data availability

Microbiota sequencing data and metabolite data will be available from the Sequence Read Archive (SRA) and the MetaboLights database for Metabolomics experiments (https://www.ebi.ac.uk/metabolights) upon request.

## 9 Declaration of interest

The authors declare no competing interests.

## 10 Acknowledgements

The Technical University of Munich provided funding for the ZIEL Institute for Food & Health, animal facility support, technical assistance and support for 16S rRNA gene amplicon sequencing. Qu Guojing provided assistance with bomb calorimetry experiments and preliminary data collection.

## Abbreviations

BA: bile acid
*Bmal1*: Brain and Muscle ARNT-Like 1
CCGs: clock-controlled genes
*Cry1*: cryptochrome circadian regulator 1
CT: circadian time
*Dbp*: D Site of Albumin Promoter (Albumin D-Box) Binding Protein
DD: constant darkness
EC: Enzyme Commission
*Ef1a*: Elongation factor 1-alpha
*Fabp2*: Fatty Acid Binding Protein 2
GF: Germ-free
GI: gastrointestinal
GUniFrac: Generalized UniFrac
*Glut2*: Glucose transporter 2
*Hdac3*: Histone Deacetylase 3
*Ifabp*: Intestinal-type fatty acid-binding protein
LD: 12 hours light and 12 hours darkness schedule
LEFSE: LDA effective score
NMR: Nuclear magnetic resonance
*Per2*: Period 2
PICRUST: Phylogenetic Investigation of Communities by Reconstruction of Unobserved States
*Pparγ*: Peroxisome Proliferator Activated Receptor Gamma
qRT-PCR: Quantitative real-time PCR
*Rev-erbα*: Nuclear receptor subfamily 1 group D member 1
SCFA: short-chain fatty acid
SCN: suprachiasmatic nucleus
SPF: specific-pathogen free
SSW: simulated shift work
T2D: type 2 diabetes
UPL: Universal Probe Library system
zOTUs: Zero-radius operational taxonomic units
ZT: *Zeitgeber* time

## 14 Supplementary Material

### 14.1 Supplementary Figure Legends

**Supplementary Figure 1** (A) Circadian profiles of relative and quantitative abundance of bacterial families in *Bmal1^SCNfl/-^* mice and their controls. (B) Heatmap illustrating the quantitative abundance of 412 zOTUs (mean relative abundance > 0.1%; prevalence > 10%). Data are ordered based on the zOTUs phase in the controls and normalized based in the peak of each zOTU. (E) Significance and amplitude (based on JTK_CYCLE) of all zOTUs (top) and phase (based on cosine regression) distribution (bottom) in both genotype, dashed line represents adj. p-value = 0.05 (JTK_CYCLE). (D) Taxonomic tree of zOTUs losing rhythmicity in *Bmal1^SCNfl/-^*mice based on relative analyses. Taxonomic ranks were indicated as phylum (outer dashed ring), then family (inner circle) and genera (middle names), each zOTU represented by individual branches. (E) Circadian profile of relative abundance of example zOTUs losing rhythmicity in *Bmal1^SCNfl/-^* mice. (F) Bar charts illustrate the alteration in abundance (adj. p-value ≤ 0.05) and fold change of zOTUs losing rhythmicity in *Bmal1^SCNfl/-^* mice. Significant rhythms according to cosine-wave regression analysis (p-value ≤ 0.05) are visualized with a solid line, while data connected by dashed line indicate arrhythmicity. n = 6 mice/time point/genotype. Data are represented as mean ± SEM.

**Supplementary Figure 2** LDA score of MetaCyc Pathways characterizing the differences between *Bmal1^SCNfl/-^* mice and their control. (B) Fecal Lactic and Valeric acid concentrations in both genotype. (C) Circadian profile of fecal SCFA. (D) Fecal bile acid concentration. Significant rhythms according to cosine-wave regression analysis (p-value ≤ 0.05) are visualized with a solid line, while data connected by dashed line indicate arrhythmicity. Mann Whitney U test was used to determine the differences between groups. n = 6 mice/time point/genotype. Data are represented as mean ± SEM. Significance * p ≤ 0.05, ** p ≤ 0.01, *** p ≤ 0.001, **** p ≤ 0.0001

**Supplementary Figure 3** (A) Summary of running wheel activity in day and night of LD and SSW group of mice. (B-C) Diurnal profile of relative abundance of major phyla (B) and family. (D-E) Heatmap illustrating the relative (D) and absolute (E) abundance of 473 zOTUs (mean relative abundance > 0.1%; prevalence > 10%). Data are ordered based on the zOTUs phase in the controls and normalized based in the peak of each zOTU. Significance and amplitude (based on JTK_CYCLE) of all zOTUs (bottom) and phase (based on cosine regression) distribution (top) in both light condition, dashed line represent JTK_CYCLE adj. p. value = 0.05. (F) Diurnal profile of example zOTUs. (G) Taxonomic tree of zOTUs losing rhythmicity in SSW mice based on relative analyses. Taxonomic ranks were indicated as phylum (outer dashed ring), then family (inner circle) and genera (middle names), each zOTU represented by individual branches. (H) Microbial composition analysis on the phyla and family level of the fecal microbiota in Lübeck and Munich. LDA score of MetaCyc Pathways characterizing the differences LD and SSW. Significant rhythms according to cosine-wave regression analysis (p-value ≤ 0.05) are visualized with a solid line, while data connected by dashed line indicate arrhythmicity. Significant phase shifts (p ≤ 0.05) are indicated with the number of hours of phase shift. Two-way ANOVA was used to assess the change in activity. n =4-5 mice/time point/genotype. Data are represented as mean ± SEM. Significance * p ≤ 0.05, ** p ≤ 0.01, *** p ≤ 0.001, **** p ≤ 0.0001

**Supplementary Figure 4 SSW has no impact on GI weight and body composition** (A) Bar charts representing colon and jejunum weight and density (B) Bar charts illustrating NMR data of SSW and control group. n = 4-5 mice/light condition. Mann Whitney U test was used to determine the differences between groups. Data are represented as mean ± SEM. Significance * p ≤ 0.05, ** p ≤ 0.01, *** p ≤ 0.001, **** p ≤ 0.0001

### 14.2 Supplementary Tables

**Supplementary Table 1: Microbial rhythmicity analysis of genetic and environmental circadian disruption mouse models:** showing microbial rhythmicity and amplitude according to JTK analysis of relative and absolute abundance of

## Notes

### Competing Interest Statement

The authors have declared no competing interest.

