## Supplementary figures and images for "Genetic and environmental circadian disruption induce metabolic impairment through changes in the gut microbiome"

### Supplemental Figure 1

##### Supplementary Figure 1

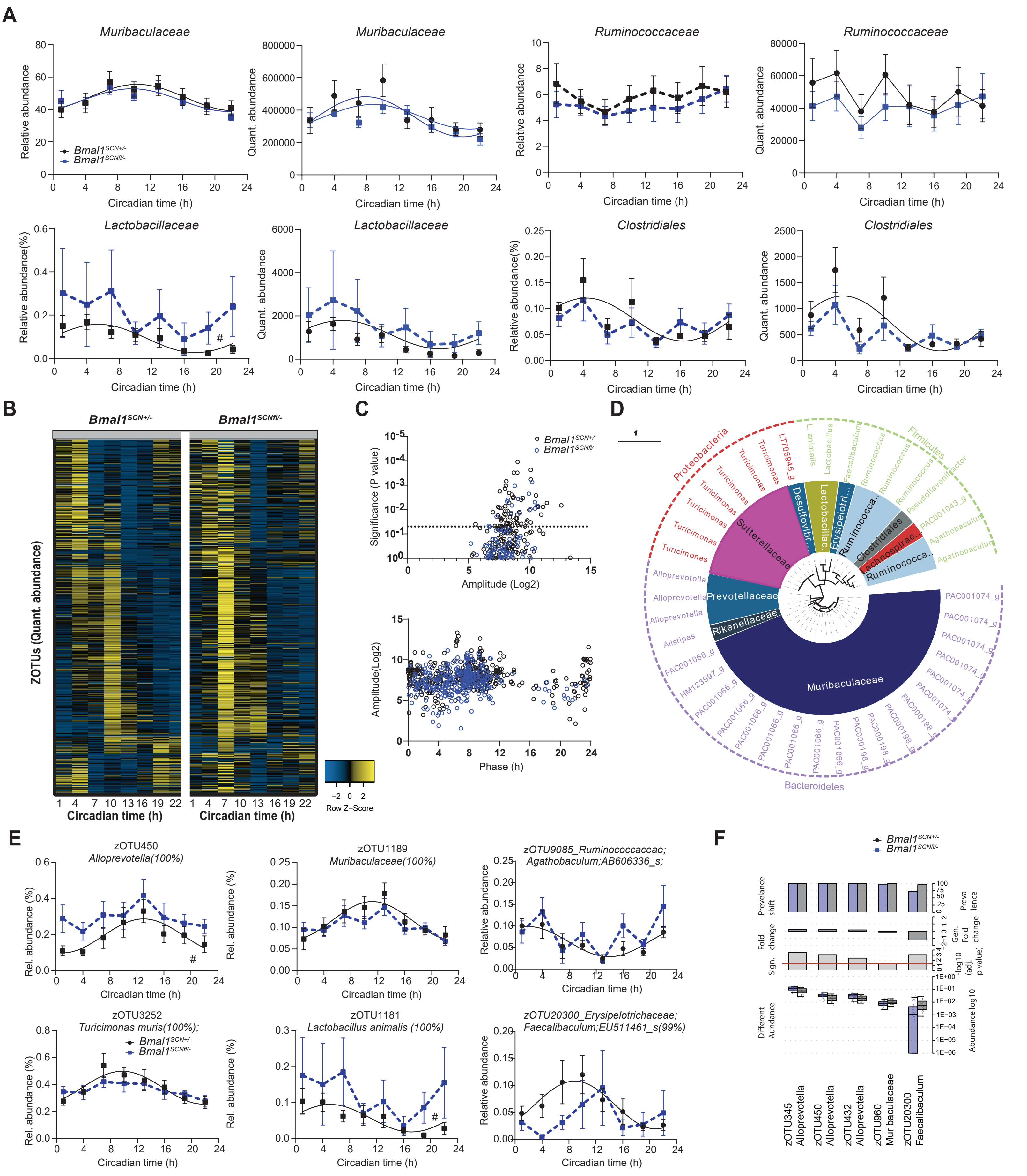

### Supplemental Figure 2

#### Supplementary Figure 2

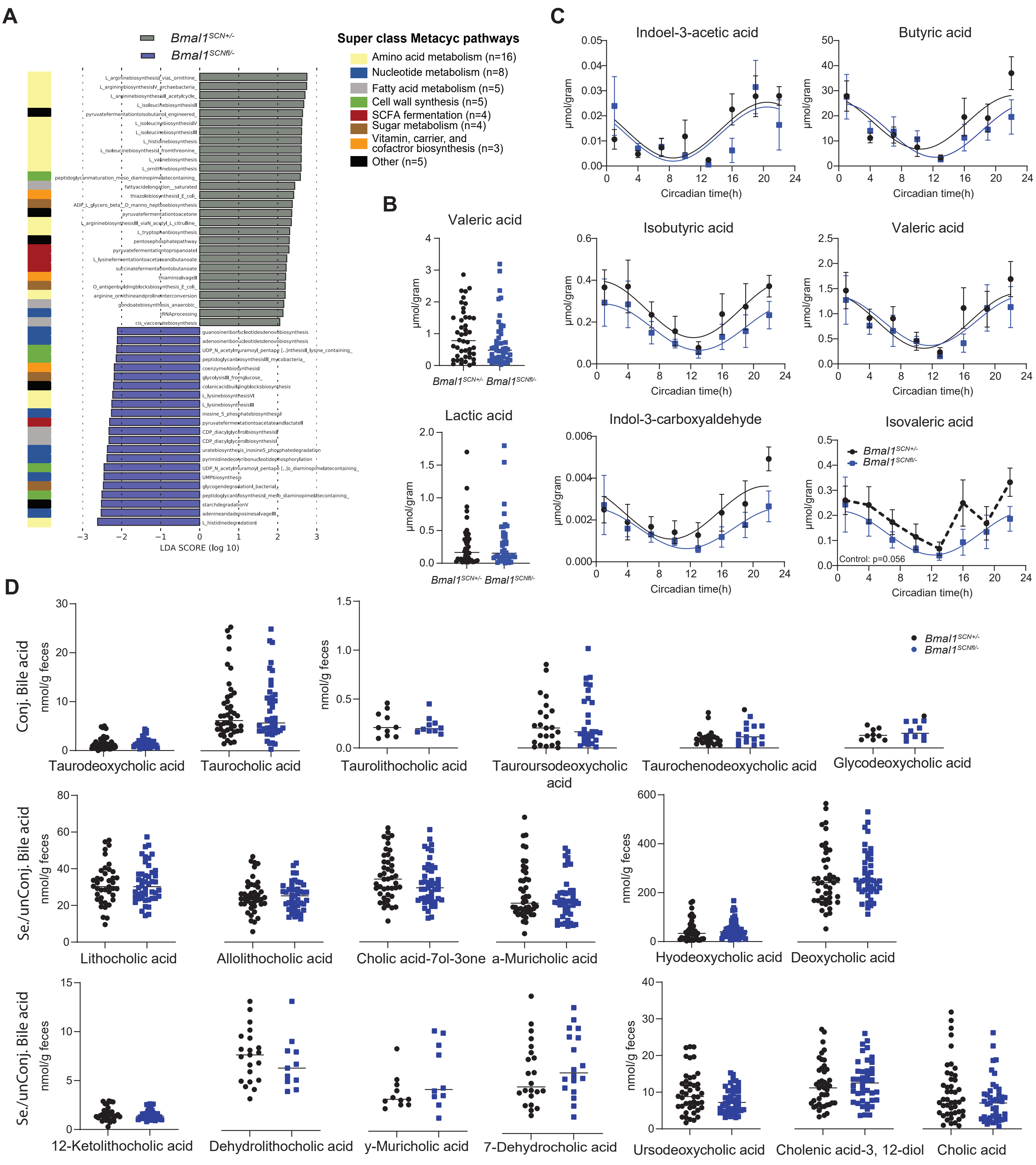

### Supplemental Figure 3

##### Supplementary Figure 3

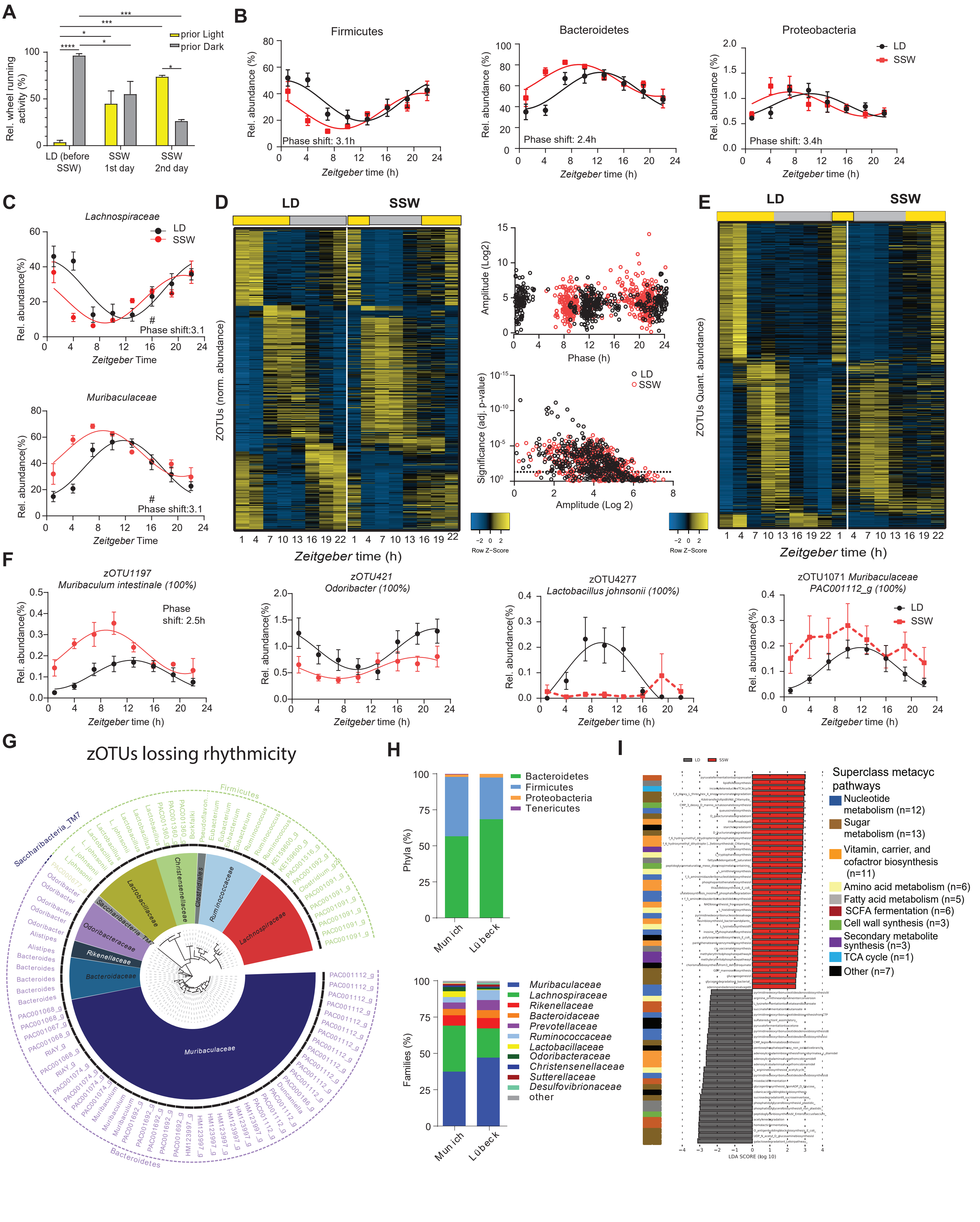
